# The history and organization of the Workshop on Population and Speciation Genomics

**DOI:** 10.1101/2022.05.14.491932

**Authors:** Julia MI Barth, Scott Handley, Daniel Kintzl, Guy Leonard, Milan Malinsky, Michael Matschiner, Britta S Meyer, Walter Salzburger, Jan Stefka, Emiliano Trucchi

**Affiliations:** Zoological Institute, Department of Environmental Sciences, University of Basel, Vesalgasse 1, 4051 Basel, Switzerland; Department of Pathology and Immunology, Washington University School of Medicine, 660 S. Euclid Ave., St. Louis, MO 63110-1010, USA; Infocentrum Český Krumlov, náměstí Svornosti 2, 38101 Český Krumlov, Czech Republic; Department of Biology, University of Oxford, 11a Mansfield Road, Oxford OX1 3SZ, UK; Institute of Ecology and Evolution, University of Bern, Baltzerstrasse 6, 3012 Bern, Switzerland; Natural History Museum, Unversity of Oslo, Sars’ gate 1, 0562 Oslo, Norway; Max Planck Institute for Evolutionary Biology, August-Thienemann-Str. 2, 24306 Plön, Germany; Institute of Parasitology, Biology Centre CAS and Faculty of Science, University of South Bohemia, Branišovská 31, 37005 České Budějovice, Czech Republic; Department of Life and Environmental Sciences, Marche Polytechnic University, Via Brecce Bianche, 60131 Ancona, Italy

## Abstract

With the advent of high-throughput genome sequencing, bioinformatics training has become essential for research in evolutionary biology and related fields. However, individual research groups are often not in the position to teach students about the most up-to-date methodology in the field. To fill this gap, extended bioinformatics courses have been developed by various institutions and provide intense training over the course of two or more weeks. Here, we describe our experience with the organization of a course in one of the longest-running extended bioinformatics series of workshops, the Evomics Workshop on Population and Speciation Genomics that takes place biennially in the UNESCO world heritage town of Český Krumlov, Czech Republic. We list the key ingredients that make this workshop successful in our view, and describe the routine for workshop organization that we have optimized over the years. We report the results of a survey conducted among past workshop participants that quantifies measures of effective teaching and provide examples of how the workshop setting has led to the cross-fertilisation of ideas and ultimately scientific progress. We expect that our account may be useful for other groups aiming to set up their own extended bioinformatics courses.

## Background

Research in organismal and evolutionary biology has been transformed over the past two decades due to the ongoing revolution in genomic technologies. The ever-increasing availability of sequencing data has ushered in a wealth of insights into evolution, but it has also brought tremendous challenges to the research community. The analysis of large genomic datasets is becoming increasingly unfeasible on personal computers and with a graphical user interface, and instead requires more and more often the use of computer servers and command-line tools[1, 2]. The rapid development of genomic technologies is accompanied by a variety of new methodological approaches, leading to a fast turnover of computer programs for evolutionary genomics. Keeping track with the latest methodological developments is therefore often a time-consuming task that can be beyond the capacity of individual research groups, especially those for whom genomics is not central to their research. In addition, the methods that group leaders learned during their PhD or postdoctoral years can be outdated by the time new students arrive and would like to be trained.

To fill this gap in cutting-edge training opportunities, several extended bioinformatics courses are being offered[3], such as those organized by the European Molecular Biology Organization (EMBO), the universities of Barcelona and Leipzig, or Evomics (Table 1). In these courses, students from different research groups and countries typically come together for two or more weeks of intense learning, free from the distractions of the regular work day. The courses comprise lectures and computer activities taught by internationally-renowned faculty that often includes the scientists who developed the methodologies that are being taught. In contrast to the teaching within research groups, this compact setting allows a large number of participants to simultaneously receive first-hand advice on the use of a collection of the latest computational tools, by some of the field’s most knowledgeable instructors. Conveniently, all participants in such a course can practice the use of bioinformatics tools with programs and example datasets that are usually preinstalled on a server infrastructure, thus avoiding potential difficulties with program installations on their own computers. Furthermore, by bringing together international participants and faculty, extended bioinformatics courses are an important opportunity for networking, among participants and between participants and faculty members.

**Table 1.**
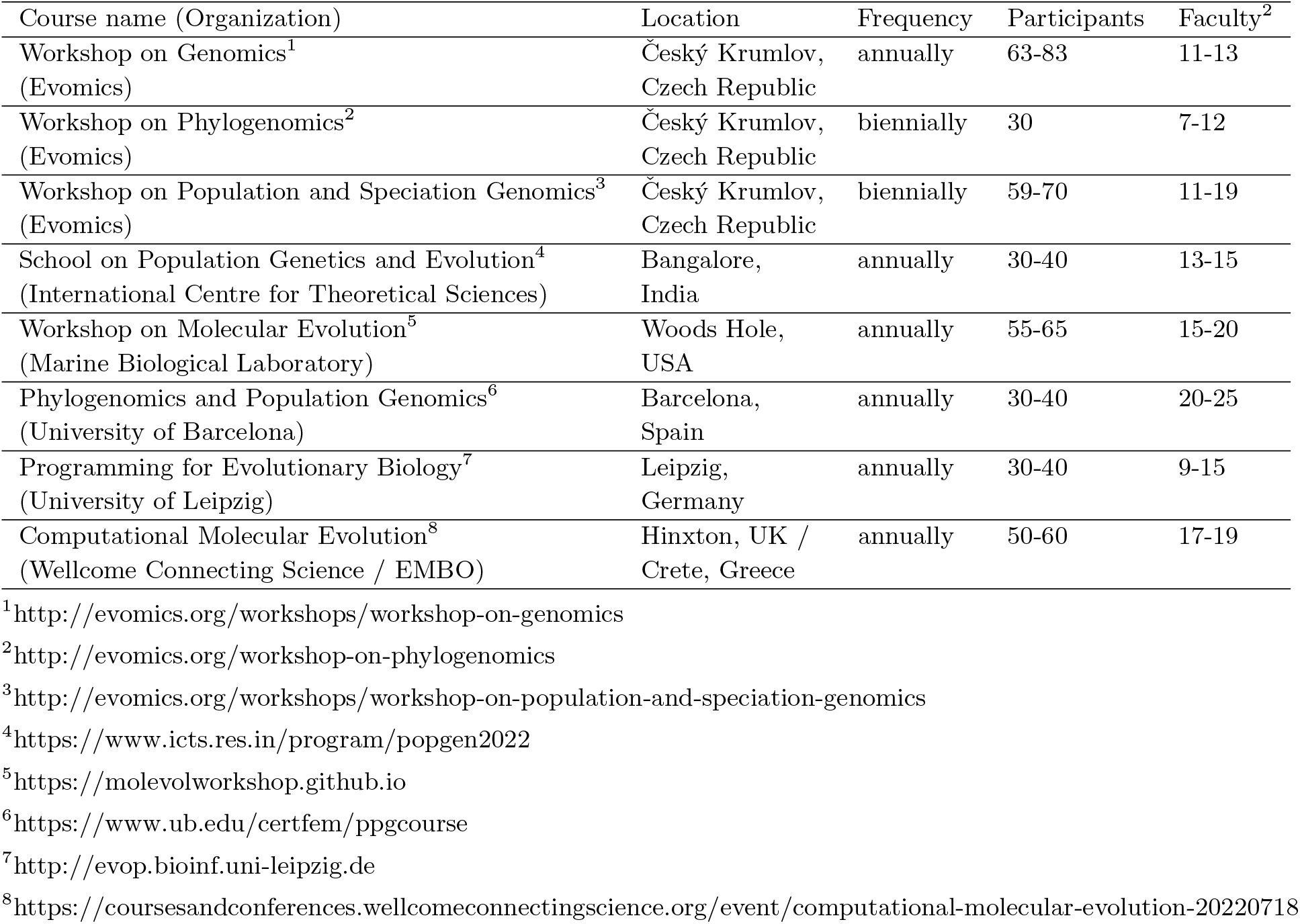
International extended bioinformatics courses in evolutionary biology. Shown are all courses with a duration of two weeks or more that focus on the use of bioinformatics applied to evolutionary questions and were advertized through Evoldir (https://evol.mcmaster.ca/evoldir.html) between 2017 and 2021. The numbers of participants and faculty members also refer to courses in these years, counting both faculty members that are part of the workshop team as well as invited faculty. Courses are sorted by organization.

We here describe the structure, the organization, and the lessons learned from one of the largest and longest-running extended bioinformatics courses, the Evomics Workshop on Population and Speciation Genomics (WPSG). This workshop first took place in 2016 in Český Krumlov, Czech Republic, and is rooted in a workshop series that began as early as 1988. All authors of this article are members of the organization team of the WPSG and and have recently organized its fourth edition. Over the past editions of the WPSG, but also building upon experience from the earlier workshop series, we identified what we consider the key ingredients of a successful workshop and established an efficient organization routine covering the year preceding the workshop. Our hope is that others can learn from our success and transpose these lessons to their own events. The demand for up-to-date scientific training to complement traditional university education has not waned in previous years and there are likely large swaths of students around the world without access to such events. We anticipate that our experience with the organization of the WPSG may be useful for other teams that take on the challenge of organizing similar events around the world in order to better serve the students’ hunger for additional high-level training opportunities.

## The deep roots of the Evomics Workshop on Population and Speciation Genomics

The WPSG arose out of a long legacy of similar training opportunities (Fig. 1). A direct chronology can be traced back to the workshop culture pioneered at the Marine Biological Laboratory (MBL) in Woods Hole, Massachusetts, USA. More specifically, the workshops organized in Český Krumlov adopted much of the underlying teaching philosophy of the Workshop on Molecular Evolution originally organized by MBL faculty since 1988 (https://www.mbl.edu/education/courses/workshop-on-molecular-evolution). This philosophy includes 1) extensive access to faculty from around the world, 2) geographic isolation to promote networking and intensive study, and 3) high-level and specific instruction not normally taught in regular university curricula. Around the mid-2s000s, it became clear that the demand for this type of training environment far exceeded the capacities available at the MBL. Thus, in order to serve more participants, members of the MBL faculty, as well as previous participants of the Workshop on Molecular Evolution, decided to make an attempt to propagate these ideas at other venues around the world.

**Fig. 1.**
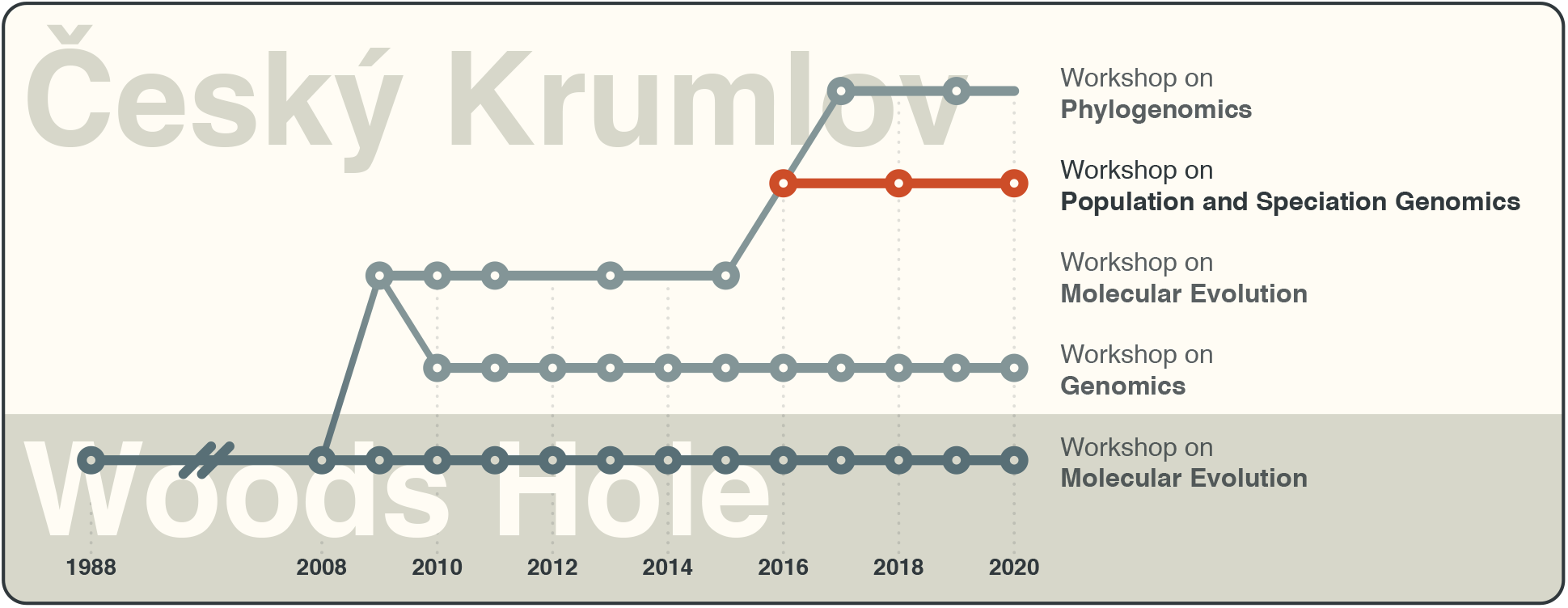
The evolution of the Workshop on Population and Speciation Genomics. Nodes indicate years in which a given workshop took place. The Woods Hole version of the Workshop on Molecular Evolution took place every year since 1988, which is indicated by the broken line between 1988 and 2008. The Workshop on Population and Speciation Genomics is highlighted in red.

While the principal teaching philosophy outlined above made up the core framework used to develop a successful workshop, there was a less quantifiable *je ne sais quoi* that those involved with a workshop at the MBL had come to know and love. This more subjective quality was reflected by the use of terms *idyllic, isolated, social*, and *experience*. There is something special that occurs when a group of people are isolated from their day-to-day business and are able to speak and work in new surroundings. The MBL at Woods Hole offered these qualities by being relatively isolated from any major city and by being a delightful venue to spend a week or two with fellow scientists. A primary task of setting up a workshop at a new location would be to adhere to the core framework, but also to identify a setting suitable of offering up the less tangible qualities that were so natural to Woods Hole.

Because of the large number of European participants and the financial and scheduling challenges many prospective participants experienced when attempting to attend the workshops in Woods Hole, a decision was made to start with an European version of the Workshop on Molecular Evolution. While Europe offers numerous idyllic and isolated locations, a key moment came with Michael Cummings, then the director of the Workshop on Molecular Evolution at Woods Hole, recalling a visit he made to Český Krumlov after giving a presentation at the nearby University of South Bohemia in České Budějovice. Michael, who had been intimately involved in developing the framework and knew the positive qualities offered by Woods Hole, immediately recognized that in Český Krumlov it may be possible to replicate and perhaps even improve upon the training model established at Woods Hole.

Following a year of preparation with the town of Český Krumlov, the first European edition of the Workshop on Molecular Evolution took place in 2009 (Fig. 1). The workshop was almost identical in faculty, topics, and schedule with that held in Woods Hole and was attended by over 100 participants from over 20 countries. The event was viewed as a resounding success in meeting an unmet demand from the scientific community and in finding a home for future workshops.

Concurrently, a sea change was occurring in the evolutionary biological sciences research community. Next-generation sequencing technologies provided by companies like 454 and Solexa were transforming the way biologists obtained and thought about genetic data. This change gave individual research groups access to millions or even billions of pieces of genetic data, capable of answering questions on an unprecedented scale. While these modern sequencing technologies were revolutionizing evolutionary research, the biological research community was not particularly well-prepared to manage and analyze the unprecedented volumes of data produced. The time of the “evolutionary biologist as a big data scientist” had arrived and the demand for training on how to handle big genomic data became very high.

As the course model for the Workshop on Molecular Evolution in both the US and Europe had proven so effective for training scientists from diverse backgrounds, it became clear that this may transfer well to other topics, including on how to manage modern sequencing data. Thus the Workshop on Genomics was born in Český Krumlov, designed to provide scientists who had little to no background in computational techniques with the tools necessary to pursue a modern sequencing project from start to finish. The same core principles of providing access to faculty from around the world in an intensive and idyllic setting were adhered to. Since the initial launch in 2010, the Workshop on Genomics has trained over 700 people from dozens of countries. It has since been running on an annual basis, only skipped for one year due to the COVID-19 pandemic. The efforts to organize the Workshop on Genomics have thus been a success, and ideas to establish similar courses on other topics began to emerge after a few years of working in Český Krumlov.

One of these ideas was to establish a world-leading effort in high-level training in the areas of population genetics and phylogenetics. Both of these research communities had already been transformed by modern sequencing technologies, and since the mid-2000s had seen the rapid development of new tools and techniques to analyze immense datasets. Therefore, the Workshop on Molecular Evolution was retired from the Český Krumlov curriculum to refocus on how to handle large datasets in a population genomic and phylogenomic context. The WPSG was developed as a new initiative designed to leverage the workshop framework established in Český Krumlov for the Workshop on Genomics and apply it to the rapidly evolving field of population genomics. Soon after, the Workshop on Phylogenomics was added to the set of workshops offered by Evomics in Český Krumlov. The establishment of these three high-level scientific training workshops has since solidified Český Krumlov as a key location for obtaining up-to-date, intensive training on a variety of topics.

## The ingredients of Evomics workshops

### The town

#### Hotels and restaurants

Despite being a UNESCO enlisted “tourist magnet” in the summer months, Český Krumlov is a relatively quiet town during the low winter season when the workshops usually take place. Yet, most hotels and restaurants remain open year-round and the prices are modest in winter. The workshop duration is a full two weeks, thus a wide selection and availability of accommodation and boarding from low-cost options, such as youth hostels and pensions, to a more up-end type of hotels is important to allow both the students on tight budgets as well as more senior participants to select lodging according to their needs. The town has plenty of choices for dining. This is preferable over catering as it provides a diverse cuisine, variation in ambience, and flexibility in group size,, and because it reduces food waste as each participant can order according to their appetite. Restaurants, cafeterias, and bars within a few minutes’ walk from the workshop venues provide access to meals and refreshments around the whole day and allow for impromptu meetings in smaller groups or “over the lunch” discussions between participants and faculty.

#### Transport

The location of the workshop in a relatively small town, isolated from the distractions of larger cities, allows for stronger connection between the participants and the faculty. Due to the relative remoteness of the location, nearest international airports are over 150 km away. However, the town is served by frequent public transport connection (railway and bus service) as well as by private “on demand” transport services, which come at an affordable price range. Assistance is provided to the faculty members with arranging transport, and detailed advice on possible options is given to the participants.

#### Local support

Given that the workshop is not hosted by an academic institution but run by organizers scattered across several continents, building up a strong relationship with the local council was essential. Particularly, the town’s tourist office (Infocentrum; https://www.ckrumlov.info/en/infocenter-cesky-krumlov/) provides the workshop team with all necessary service. The office’s local contact point, Daniel Kintzl, has become an indispensable part of the team, providing assistance with facilities’ rental agreements, with everyday *ad hoc* or even emergency (e.g., hospital) requirements, and by organizing leisure activities, like yoga classes, town sightseeing, and wine tasting.

### The facilities

#### Lecture and computer activity venues

Český Krumlov offers several venues that meet the requirements of a heavily computation-oriented workshop with over 60 participants. Topical lectures take place in the town’s theatre hall, which offers a representative venue with necessary audio and video equipment for lectures, and a cafeteria for coffee-break refreshments. A large room arranged for computer activities is located in a nearby building (the medieval House of Prelate), which is equipped with fast wired and wireless internet.

#### Computational resources

A big aspect of the organisation of the workshops is the compute provision. For this we rely on several systems running smoothly together: 1) participant laptops, 2) a cloud based virtual machine service, and 3) the WiFi and internet connections specially provisioned each year in the House of Prelate and other venues.

Each participant is required to bring a modern laptop, which they use throughout the workshop. However, the majority of any computer activity is not performed locally on their machines, but instead on virtual machines based in the cloud. All what is required from each participant to use these virtual machines is a “terminal” program for running command-line interface software and a modern web browser to run graphical-user-interface programs. To allow the latter, the use of Apache Guacamole (https://guacamole.apache.org) – a web server providing access to cloud-based desktops – has proven effective. This setup solves many problems: First, dedicated software does not need to be installed on participants’ laptops, which can be difficult on those that are administered by the IT departments of the participants’ universities. Second, input data do not need to be downloaded, which could be slow and consume valuable teaching time as some activities require gigabytes of data (note that all participants use the same internet connection). Third, computer activities do not depend on the speed of the participants’ laptops, the operating systems installed on these laptops, or their keyboard layouts.

In earlier editions of Evomics workshops, virtual machine images were provided to each participant on USB drives so that they could run these on their laptops. However, it was a costly and slow process to prepare over 60 USB drives before the commencement of the course and almost impossible to correct any errors in materials, data, or software. To overcome these issues, Evomics workshops now rely on the cloud systems of Amazon Web Service (alternatives include Google Cloud, Microsoft Azure, and XSEDE) that are easy to access and administer, and provide the ability to build virtual machines – Amazon Machine Images (AMI) – via the Amazon Elastic Compute Cloud (EC2). The virtual machines can then be launched as multiple instances, producing an identical virtual machine instance for each participant.

The development of AMIs for Evomics workshops begins with a base image containing the latest version of Ubuntu with a set of standard bioinformatics software tools and workshop branding; the software ANSIBLE (https://www.ansible.com) has proven particularly useful for automating this process. Then, before each workshop, we ask the faculty to provide a list of specific software and data that will be needed for their tutorials. These are then built directly into the AMI and tested the week before the workshop commences, to allow for any bugs, issues, or software incompatibilities to be fixed in time.

The main advantage of this process is speed, as only one image needs to be kept up-to-date and any changes can be distributed near-instantaneously to the participants. Workshop participants are able to continue working on their specific instance throughout the course. The running instances can be easily administered by the workshop team via Amazon’s console; allowing us to check for current costs, any misuse or security issues, and to turn them on and off between sessions (this reduces the workshop’s costs, as, for example, 60 “t2.medium” instances running for nine hours per day, with 500 GB Elastic Block Storage, cost around 1,500 USD for the duration of the workshop). Importantly, the computational resources of AMIs (e.g., CPUs and memory) can be flexibly changed to adapt to the requirements of each activity.

To allow keen participants to continue computer activities after their departure, the workshop’s AMIs usually remain available for one year. However, participants using AMIs after the workshop will need to pay for their own compute time.

#### Website

The central platform for sharing of workshop information is the Evomics website (https://evomics.org). The website houses all of the training materials provided by each workshop edition in perpetuity, thus minimizing the concerns of many participants to finish all activities during their short stay in Český Krumlov. Participants are always welcome to continue their work and share the knowledge they gained after the return to their home institutions. During the workshop, the website becomes a communication center for document sharing, announcing schedule/venue updates, and providing background on faculty and other course participants. Announcements about future workshops are also made through the workshop website. The website is therefore a living document for documenting historic, present, and future workshop information.

#### Outreach and internal communication

To increase visibility among scientists and to announce registration deadlines, the workshop team promotes the WPSG on various social media platforms (e.g., Twitter and Facebook) as well as email lists for evolutionary biologists (e.g., Evoldir; https://evol.mcmaster.ca/evoldir.html). To ensure good management of the various tasks before and during the workshop, much of the communication within the team takes place in a shared online workspace, such as Slack (https://slack.com). This allows the team to communicate on specific topics, to discuss in subgroups, and exchange direct messages between each other. Besides the online workspace used by the workshop team for organization, another workspace is created during the workshop and used by the participants.

### The teaching

#### Co-directors

At the core of the workshop organization is a group of eight co-directors, who are all active scientists in research fields closely related to the workshop’s main topics. Besides having a general research interest in evolutionary biology, population genomics and phylogenomics, they are also familiar with hands-on work on bioinformatic analyses and some of them implement novel methodological approaches to analyze genomic data. The range of academic positions held by the co-directors is between senior postdocs and professors.

#### Faculty

The faculty line-up is selected for every workshop edition from a long list of international researchers. The list is compiled on the basis of 1) recent publication of relevant bioinformatic tools and methodological approaches in population and speciation genomics, 2) adequate representation of the different topics planned in the workshop agenda, 3) diversity in gender and seniority, 4) previous workshop attendance as faculty, teaching assistant, or participant, 5) information available on teaching and interaction skills. All faculty members are invited to stay in Český Krumlov for the whole workshop duration, to maximize the potential for interaction with participants also outside of their lecture, computer activity, and faculty lunch (see below).

#### Teaching assistants

Teaching assistants (TAs) are an important component of the workshop team. They have usually taken the workshop as participants before and approached the co-directors to become TA in a future workshop. The TAs are typically senior PhD students or junior postdocs with an interest and first-hand experience on many of the topics and bioinformatic analyses covered in the workshop. Together with co-directors, TAs support faculty members during the different computer activities by answering questions from participants and helping to solve computer issues, and by testing the computer activities beforehand. Usually, between two and three TAs are employed, depending on the number of admitted participants.

#### Preparation material

Both faculty members and co-directors are asked to suggest relevant didactic material such as articles, text books, online presentations, and tutorials to be made available to the participants a few weeks before the workshop as preparatory material on the workshop website. This material is meant to bring the participants to the same minimal starting level (e.g., through UNIX tutorials), to refresh basic statistics and population genetics concepts, and to prepare the ground for the topics and bioinformatic analyses which are presented during the workshop lectures and computer activities. The preparatory material is updated and adjusted on the basis of the faculty lineup before each workshop edition.

#### Supporting teaching material

The actual teaching material consists of lecture presentations and computer activity tutorials. The presentation and tutorial materials are made available on the workshop web page on the day when the lectures and computer activities take place, and they remain permanently and freely available, unless a faculty member objects to this Open Access policy. All faculty members in charge of computer activities are asked to provide their tutorials at least one to two weeks in advance, so that these can be thoroughly tested on the workshop’s virtual machine by co-directors and TAs.

### The mentoring

#### Faculty lunches

Workshop participants are invited to join so-called “Faculty Lunches”, that is, opportunities for discussion that are offered daily with a different faculty member. This platform is explicitly designed for those participants who are interested in a dialogue-oriented event in a small group setting. Registration for faculty lunches is done through an online time management tool to keep track of the number of participants, and is limited to two faculty lunches per attendee. Restaurants are chosen as the location of the event, depending on the group size and subject to space availability. In some cases, the discussion is accompanied by a TA. The informal meetings have full freedom of choice of topics and are intended to facilitate interaction between participants and faculty members, increase group unity, and encourage scientific discussions.

#### Open labs

At least once during the workshop we offer the possibility to work in “Open Labs”. In these time slots, remaining questions from previous computer activities can be answered and worked on together. In addition, the open labs are often used to discuss topics according to the needs of the participants in small groups, including the co-directors and/or faculty members. These small focused groups are then spatially distributed to discuss and study in a concentrated manner. Alternatively, these time slots can also be used simply to relax or get to know the work of fellow scientists over a coffee.

### Social aspects

In addition to scientific education, a core mission and desire of the workshop is to create a social sense of community and an unforgettable memory. To this end, a number of successful events and ideas have been established over the years and continue as workshop traditions.

#### Welcome reception

Upon the arrival of the participants on the first evening, a welcome party with buffet is organized in one of the local hotels. At this reception, co-directors and TAs greet the arriving participants, start informal discussions about the course, and provide participants with information about the premises of the lecture the next morning. The evening with food and drinks therefore serves as an icebreaker to get to know each other but also to clarify initial questions and uncertainties. After the reception, the group often moves to a neighboring bar to deepen these first conversations and acquaintances.

#### Mid-workshop and farewell parties

In the middle of the workshop there is traditionally a party organized by the TAs. Of course, like any party, this event is entirely for fun and relaxation, allowing all participants to take a break from science. This offer is generally gladly accepted by a large part of the participants. Towards the end of the workshop, some keen participants then take over the organization of the farewell dinner and subsequent celebrations, which take place on the evening of the last workshop day.

#### T-shirt competition

To reinforce the workshop brand and sense of unity, participants and TAs are encouraged to create the design for a workshop T-shirt. The T-shirts are printed before the end of the workshop, allowing the participants to return home with a unique souvenir.

#### Bingo

To encourage the participants to explore and get to know the town, the TAs have developed a game in which certain places or tasks in Český Krumlov have to be found or accomplished. These tasks are arranged in a “bingo” field and the goal is to complete as many as possible within the duration of the stay. The first participant to complete the bingo wins a small prize.

#### After-work-gatherings

A collaborative decision is made each evening about where most of the workshop group will get together after the last computer activity, so that participants and faculty have a place to meet for socializing.

#### Other activities

Last but not least, there are also leisure activities that are organized by the tourist office and can be booked there. These include various sport activities (such as yoga, archery, or hiking) and cultural events (such as guided town tours, visits of the Český Krumlov castle, and wine tastings). Running, soccer, or climbing groups have also organized themselves and met during free times, thus providing the necessary balance to the intense workshop schedule.

#### Group composition

The WPSG has so far always attracted more applications than can be accepted due to space limitations, and thus, a selection of participants is required. The selection is based primarily on an estimate of the degree to which participants might benefit from attending the workshop (e.g., whether they may immediately apply knowledge from the workshop to their own data). However, attention is also paid to balancing gender, nationality, and (academic) age of the participants (Table 2). In our experience, the resulting heterogeneous composition creates a stimulating environment and productive scientific exchange.

**Table 2.**
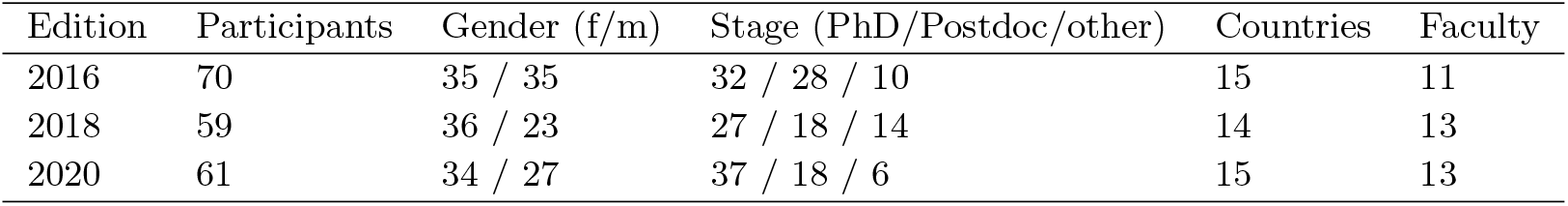
Past editions of the Workshop on Population and Speciation Genomics.

| Edition | Participants | Gender (f/m) | Stage (PhD/Postdoc/other) | Countries | Faculty |
| --- | --- | --- | --- | --- | --- |
| 2016 | 70 | 35 / 35 | 32 / 28 / 10 | 15 | 11 |
| 2018 | 59 | 36 / 23 | 27 / 18 / 14 | 14 | 13 |
| 2020 | 61 | 34 / 27 | 37 / 18 / 6 | 15 | 13 |

## Workshop preparation

Preparations for the workshop take place over the course of nearly one entire year (Fig. 2). In January in the year before a new workshop edition, the group of co-directors comes together and decides upon realization of the workshop, followed by decisions on initial tasks such as setting up and updating workshop web pages, the reservation of workshop venues, and the approximate targeted participant number. Yet, the most time-consuming task in this early organizational phase is the compilation of the new teaching schedule and faculty. Having a diverse workshop team including experts from different scientific disciplines is of great advantage, as up-to-date knowledge about the latest methodological developments as well as broad networks to leading scientists in the respective fields are indispensable to fulfill the workshop’s teaching philosophy. Upon deciding on an ideal schedule and core faculty members, the lecturers are invited and the schedule is continuously updated depending on acceptance of the invitations, taking into account the temporal availability of faculty. The aim is to finalize the schedule, officially announce the workshop, and open the registration for participants before the start of the European summer holidays in July.

**Fig. 2.**
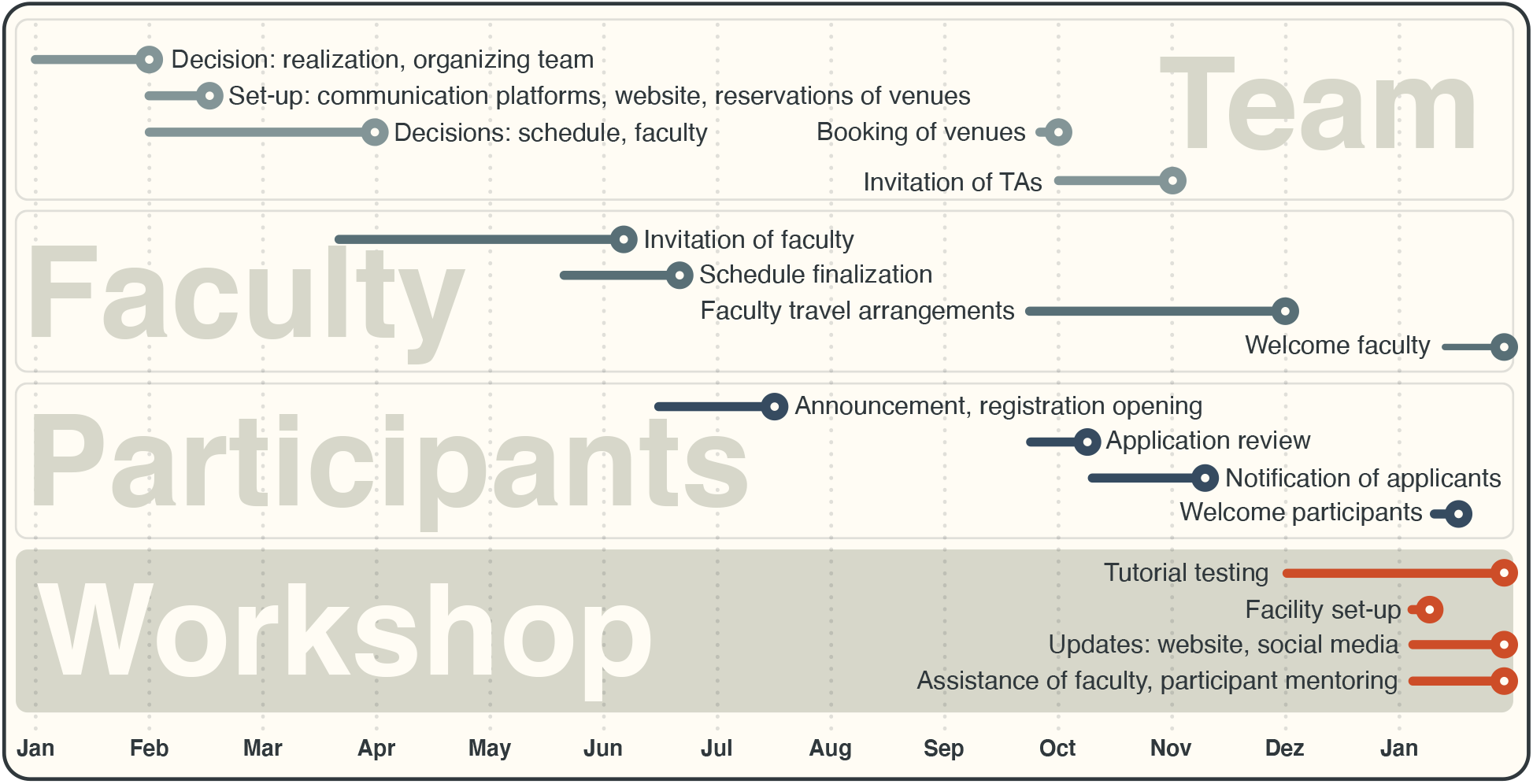
Timeline for the preparation of the Workshop on Population and Speciation Genomics. The workshop is prepared over the course of the year preceding a workshop edition, starting with the formation of the workshop agenda by the team, and followed by the recruitment of faculty, the announcement, and the selection of participants.

Registration usually remains open until mid-September, which is the latest time-point to assess whether the number of potential participants would be sufficient to hold the workshop. Upon the fulfilment of this requirement, several tasks are initiated more or less simultaneously: participant applications are reviewed and participants are notified, TAs are invited, the workshop venues are booked, the faculty members are informed about travel details and asked to confirm or book their transport and, finally, hotel rooms are booked for the workshop team and the faculty. The goal is to complete all these tasks before the Christmas break.

Before the actual workshop takes place in January/February of the following year, the faculty members are asked to provide information about required software, as well as tutorials for their computer activities, so that this information can be added to the teaching material on the workshop website, and the software and tutorials can be uploaded to the cloud servers and tested on the virtual machines. Although the workshop team begins to test the first tutorials already in December, the bulk of this task is taken care of during the preparation week at the workshop venue just before the workshop starts. The workshop-preparation cycle ends with the end of a workshop and starts again nearly exactly a year later for the next workshop.

## Measures of effective teaching

The most important metric of the workshop’s success is the high number of applications for participation that we keep receiving: for each edition of the workshop, we have been able to accept between 60 and 70 participants out of 90 to 120 applications.

Surveys among participants are conducted soon after the workshop, to understand if the workshop met their expectations and to obtain a timely evaluation of both the scientific content (lectures and computer activities) and the overall organization. Results of these anonymous surveys are in general extremely positive while, on the other hand, they have helped us to continue improving the workshop.

During the writing of this article, we asked former participants, from all past editions of the WPSG, to answer five short questions for an assessment of the long-term effectiveness of the workshop teaching and interactions. In particular, we asked whether: 1) the scientific content they were trained in during the workshop was useful for their own project’s implementation, 2) the workshop’s computer activities had been helpful to analyze their data after the workshop, 3) the network they could establish during the workshop with the other participants and faculty members was fruitful in terms of collaborations and/or career opportunities, 4) the methods and techniques acquired during the workshop were helpful to find a new position or fellowship, and if 5) they have recommended the workshop to colleagues or fellow students. Answers from 68 participants, representing a wide range of academic positions and countries of affiliation, show highly positive responses to question 1, 2, and 3, while slightly above neutral answers to question 4 (Fig. 3). At any rate, almost all of the respondents had strongly recommended the WPSG to other colleagues and students (Question 5; Fig. 3), suggesting that they appreciated the high level of teaching provided.

**Fig. 3.**
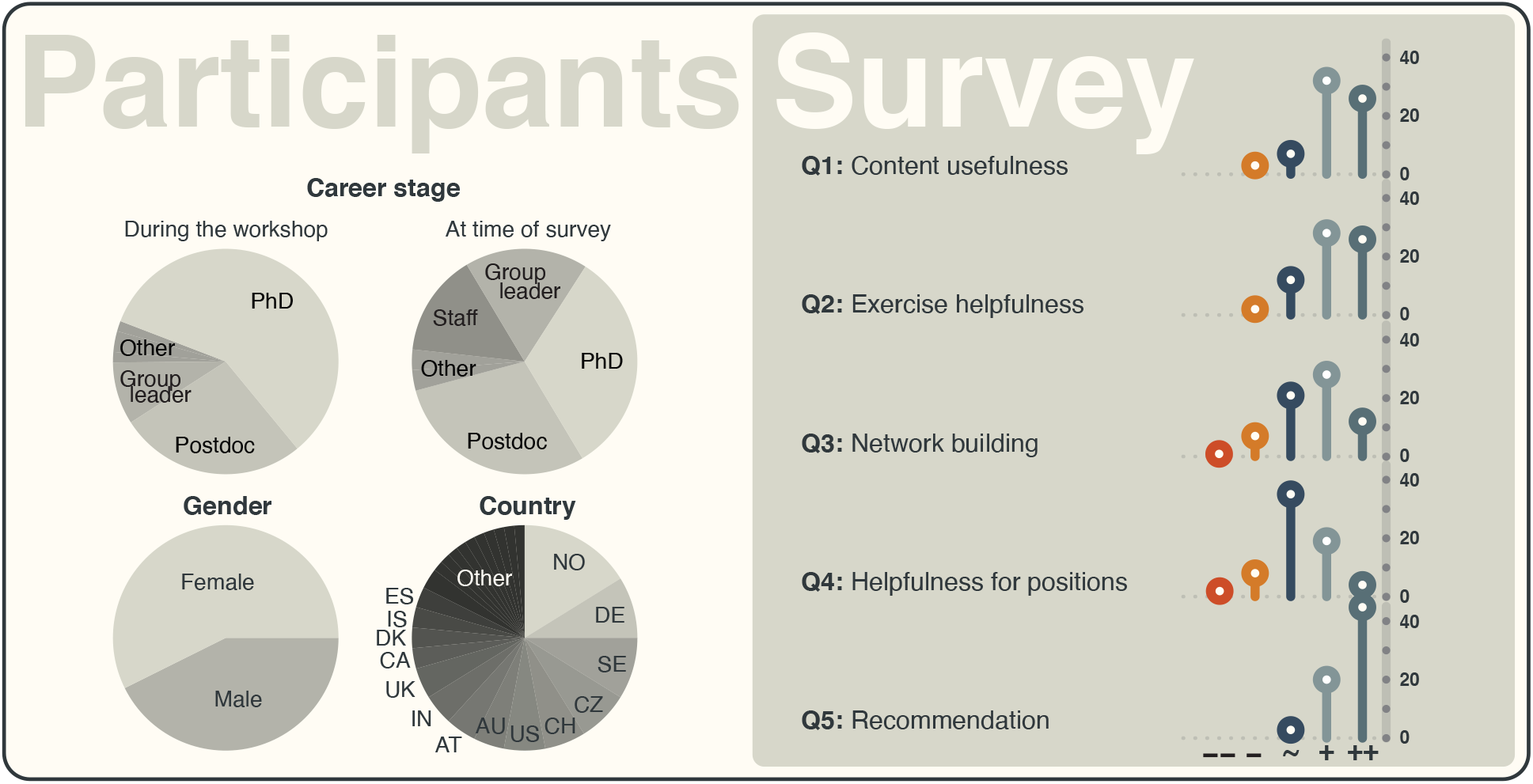
Results of a survey on workshop quality among past participants. Past workshop participants were contacted and asked five questions on the usefulness of workshop content, the helpfulness of computer activities, the effectiveness of network building at the workshop, the helpfulness of the workshop for their career, and whether they have recommended the workshop to colleagues (see main text).

## Cross-fertilisation of ideas

The unique mix of ingredients brought together at the workshop is not only beneficial to the education of participants, but also fosters a creative environment where scientific ideas mix and flourish. Participants across the breadth of the discipline share their challenges with world-class faculty and the workshop team, pointing to common problems and issues to be addressed in order to move the entire field forward. Within the workshop setting, all involved are more likely to find the focus, expertise, and the time to meet these challenges.

The benefits of such cross-fertilisation of ideas are exemplified by the fineRADstructure[4] software package, which owes its existence to the 2016 edition of the WPSG (see Box 1).

## Conclusion

With the continuous and rapid turnover of state-of-the-art bioinformatic methodology, centralized teaching of computational approaches and their application is now more relevant than ever before. The WPSG is dedicated to this type of teaching, based on lectures and computer activities by internationally renowned faculty members. As evidenced by the result of our survey among past workshop participants, the WPSG has proven effective, with a positive influence on the careers of those who attended it. However, the WPSG is not able to satisfy the educational needs of all students in evolutionary biology, due to its limited capacity. We would therefore welcome the initiation of additional extended bioinformatics courses, and aimed in this article to provide guidance by describing what we consider the most important ingredients of such a course. We regularly have participants who travel long distances to attend the WPSG (e.g., from Chile or Australia – only six nations have been more represented than Australia; Fig. 3), which implies a lack of educational opportunities closer to the home places of these participants. The spawning of new and globally distributed extended bioinformatics courses would therefore not only render bioinformatics education cheaper and more easily accessible for world-wide researchers, but would also reduce the strain on the environment caused by long-distance traveling. We hope that this article may serve to facilitate such developments.

### Box 1

**fineRADstructure**

In 2016, fineSTRUCTURE[5] was one of the premier tools for understanding population structuring and used extensively in human genetics. One of its key authors, Daniel Falush, was among the faculty at the inaugural WPSG. In interacting with participants at the workshop, it soon became apparent that the application of his tool to all the species studied by evolutionary biologists would not be possible because of the differences in genetic data used – while human geneticists typically used high quality whole genome data, many other evolutionary biologists used so-called restriction-site associated DNA (RAD) sequencing markers.

With such feedback from participants, Daniel enlisted the help of workshop team members, Emiliano Trucchi for his in-depth expertise in RAD data and Milan Malinsky for his programming and software development capabilities. The first versions of fineRADstructure, a variant of fineSTRUCTURE specifically for RAD data, were developed during interactions directly at the workshop, with a preprint and initial code released later in 2016. Today, fineRADstructure ranks among commonly used tools in evolutionary genomics and the manuscript describing the software[4] has at the time of writing accumulated more than 170 citations.

## Author’s contributions

All authors contributed to the writing of this manuscript.

## Acknowledgements

We sincerely thank the town of Český Krumlov, and particularly the employees of the Tourist Information, the Zlaty Andel Hotel, the Town Theater, the House of Prelate, and the Egon Schiele Café for being such formidable hosts for our workshop. We acknowledge financial support from the University of South Bohemia in České Budějovice that allowed us to lower workshop fees for participants. We would also like to express our gratitude to the key figures responsible for the establishment of Evomics workshops in Europe: Michael Cummings, Naiara Rodríguez-Ezpeleta, Dave Swofford, and Adam Bazinet. Without them, the WPSG would never have existed. Finally, we greatly thank all faculty members, teachings assistants, and participants who ever joined a WPSG edition (particularly those participants who answered our survey) for making these workshops the unique events that they are.

